# Model colibactins exhibit human cell genotoxicity in the absence of host bacteria

**DOI:** 10.1101/431098

**Authors:** Emilee E. Shine, Mengzhao Xue, Jaymin R. Patel, Alan R. Healy, Yulia V. Surovtseva, Seth B. Herzon, Jason M. Crawford

**Affiliations:** Department of Microbial Pathogenesis, Yale School of Medicine, New Haven, Connecticut 06536, United States; Chemical Biology Institute, Yale University, West Haven, Connecticut 06516, United States; Department of Chemistry, Yale University, New Haven, Connecticut 06520, United States; Department of Molecular, Cellular, and Developmental Biology, Yale University, New Haven, Connecticut, 06516, United States; Yale Center for Molecular Discovery, West Haven, Connecticut 06516, United States; Department of Pharmacology, Yale School of Medicine, New Haven, Connecticut 06520, United States

## Abstract

Colibactins are genotoxic secondary metabolites produced in select Enterobacteriaceae, which induce downstream DNA double-strand breaks (DSBs) in human cell lines and are thought to promote the formation of colorectal tumors. Although key structural and functional features of colibactins have been elucidated, the full molecular mechanisms regulating these phenotypes remain unknown. Here, we demonstrate that free model colibactins induce DSBs in human cell cultures and do not require delivery by host bacteria. Through domain-targeted editing, we demonstrate that a subset of native colibactins generated from observed module skipping in the nonribosomal peptide synthetase-polyketide synthase (NRPS-PKS) biosynthetic assembly line share DNA alkylation phenotypes with the model colibactins *in vitro*. However, module skipping eliminates the strong DNA interstrand cross-links formed by the wildtype pathway in cell culture. This product diversification during the modular NRPS-PKS biosynthesis produces a family of metabolites with varying observed mechanisms of action – DNA alkylation versus crosslinking – in cell culture. The presence of membranes separating human cells from model colibactins attenuated genotoxicity, suggesting that membrane diffusion limits colibactin activity and could account for the observed bacteria-human cell-to-cell contact phenotype. Additionally, extracellular supplementation of the colibactin resistance protein ClbS was able to intercept colibactins in an *E. coli*-human cell transient infection model. Our studies demonstrate that free model colibactins recapitulate cellular phenotypes associated with module-skipped products in the native colibactin pathway and define specific protein domains that are required for efficient DNA interstrand crosslinking in the native pathway.

---

Perturbations in community structure of the gut microbiota coupled with shifts in host–microbe metabolism can lead to a variety of altered host physiologies associated with diseases, such as inflammatory bowel disease and cancer.^*1–3*^ A prominent, yet rare, example of a one-pathway, one-phenotype correlation in the microbiome is that of the colibactin pathway.^*4–8*^ The colibactin gene cluster (*clb*) contains 19 genes (*clbA*–*clbS*) that encodes the biosynthesis of hybrid polyketide synthase– nonribosomal peptide synthetase (PKS–NRPS) secondary metabolites known as colibactins. The *clb* locus is found in many strains of Enterobacteriaceae including select strains of gut-commensal and extraintestinal pathogenic *E. coli* (ExPEC) and *Klebsiella pneumoniae*, among others.^*9, 10*^ The pathway is associated with virulence^*11*^ and is significantly more prevalent in patients with inflammatory bowel disease (IBD), colorectal cancer (CRC), and familial adenomatous polyposis (FAB).^*12–14*^

A growing number of studies support a causative role for the metabolites in colorectal cancer formation. For example, when co-cultured with colibactin-expressing strains, mammalian cells accumulate DNA double-strand breaks (DSBs), activate ataxia-telangiectasia mutated kinase (ATM) signaling, and undergo cell cycle arrest and senescence.^*9, 15, 16*^ *Clb+ E. coli* induce tumor formation in three *in vivo* models of CRC^*12, 17, 18*^. Synthetic colibactin derivatives of metabolites from *clb+* bacteria are genotoxic *in vitro*^*19*^. Finally, it was recently shown that incubation of exogenous DNA with colibactin-expressing strains induces DNA interstrand crosslinks in a pathway-dependent manner.^*20*^

Characterization of colibactins has been difficult; to date, no colibactin has been directly isolated from any producing strain. This has been attributed to their low levels of natural production and/or putative instability^*21*^. Progress has been made using a combination of biochemical characterization of pathway enzymes^*22–27*^, comparative metabolomics^*21, 28, 29*^, isolation from large-scale fermentation of genetically-modified mutants,^*30, 31*^ and chemical synthesis.^*7, 19, 29, 32*^ This body of research has revealed important aspects of colibactin biosynthesis, structure, and function. First, colibactins are assembled by the NRPS–PKS biosynthetic machinery as prodrugs with an appended *N*-acyl-D-asparagine side chain.^*22, 28, 33*^ These “precolibactins” are transported into the periplasm by the 12-transmembrane transporter ClbM.^*34*^ The *N*-acyl-D-Asn residue is then removed by the pathway-dedicated membrane-bound peptidase (ClbP).^*35, 36*^ Upon deacylation, spontaneous cyclization reactions occur to generate genotoxic colibactin structures.^*7, 19, 29*^

Deletion of any one of the biosynthetic genes within the colibactin pathway results in abrogation of DSBs in cell culture.^*9*^ Access to fully-functionalized colibactins is desirable to understand their trafficking from the periplasmic space and mode of action in eukaryotic cells. However, to the best of our knowledge, a metabolite that accounts for all of the genes in the *clb* gene cluster has not been isolated or predicted with experimental support.

Model (pre)colibactins alkylate and crosslink DNA *in vitro*.^*19,21*^ Previous studies identified colibactin **1**, which was detected in ClbP-proficient strains (Fig. 1A).^*29*^ Derivatives of **1**, such as the *N*-methylamide **2** and dimethylethylene diamine derivative **3** alkylated DNA *in vitro* by nucleotide addition to the electron deficient cyclopropyl residue.^*19*^

**Figure 1.**
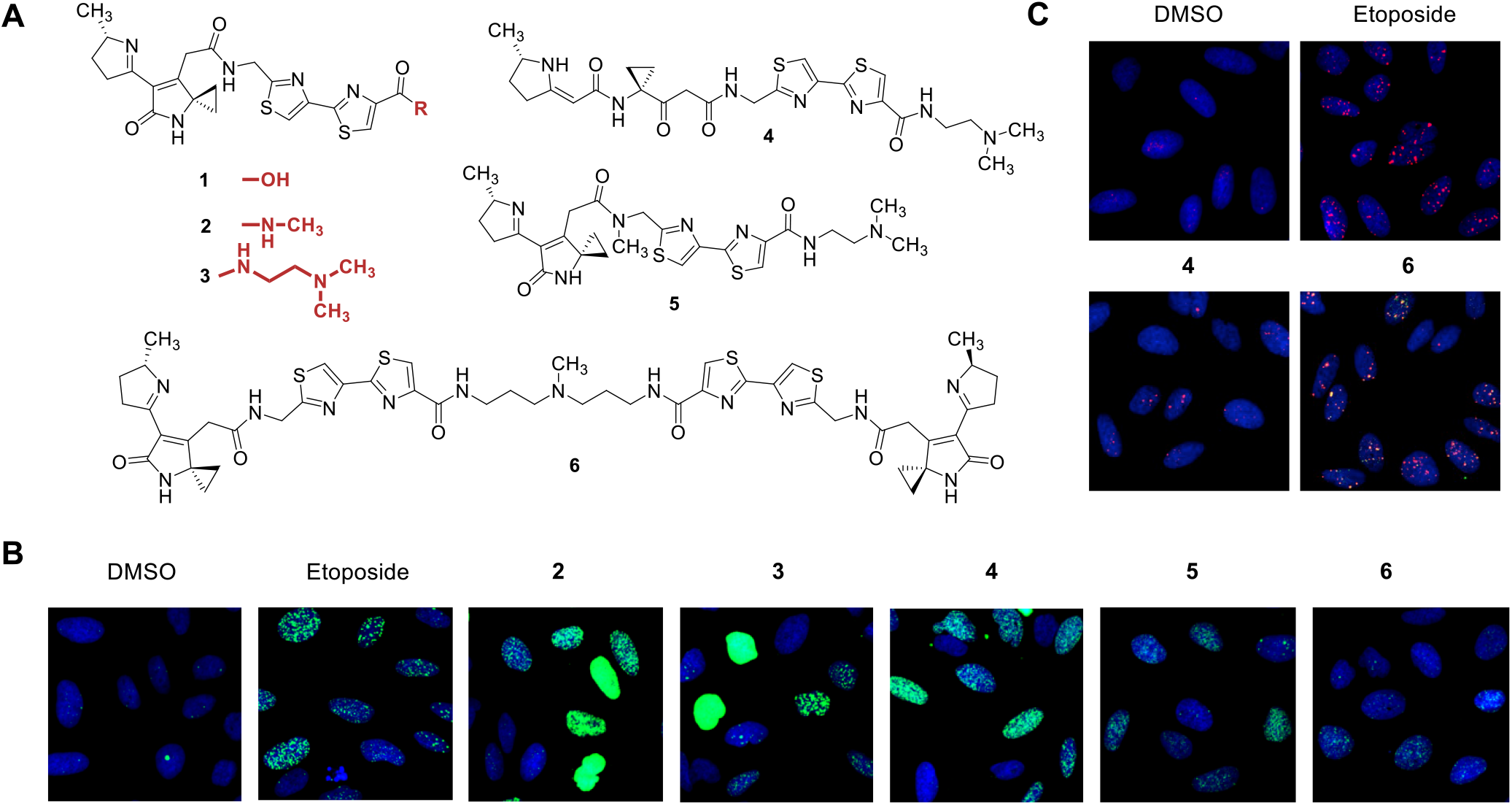
Synthetic colibactins exhibit cellular genotoxicity. (A). Structures of synthetic colibactins that previously demonstrated *in vitro* DNA alkylation activity. (B). HeLa cells were treated with 11 µm of compounds **2–6** for 4 hours and subsequently fixed and stained for a marker of cellular DNA damage, γH2AX. As a positive control, 1 µM of etoposide is shown for comparison. Quantification of γH2AX induction relative to that of etoposide is shown in the supplementary information. (C). Of the model colibactins, only dimer **6** shows significant activation of 53BP1, a downstream effector of ATR kinase signaling, a phenotype induced by the native colibactin pathway.

We have shown that linear biosynthetic products are offloaded from the pathway and that they undergo spontaneous cyclization to unsaturated imines, such as **1**^***5***^, via intermediates resembling **4**^***19***^. We previously synthesized **4** and found that it is genotoxic *in vitro*, indicating that cyclization to an unsaturated imine is facile. The *N*-methylamide analog **5**, which was designed to block formation of an inactive pyridone isomer, was also genotoxic. Finally, the dimer **6** was synthesized and examined for functional comparison. With the exception of **6**, all compounds incubated with linearized plasmid demonstrated extensive alkylation and degradation of duplex DNA, with cyclic imine and cationic cap moieties enhancing potency down to the nanomolar range. Compound **6** crosslinked DNA, as expected.^*19*^

Herein, we evaluated the abilities of the model colibactins **2**–**6** to recapitulate the genotoxic phenotypes associated with colibactins produced in transiently infected human tissue cultures. Human cell (HeLa, U2OS) DNA damage was quantified by immunofluorescence imaging of the DNA damage marker phospho-SER139-histone H2AX (γH2AX).^*37*^ The effects of the compounds incubated for 4 h at 16 doses ranging from 230 nM to 100 µM were quantified relative to 1 µM etoposide as a positive control (set at 100% effect) and DMSO vehicle as a negative control (0% effect). All of the compounds induced γH2AX activation and were cytotoxic. Aside from a few exceptions, we could establish reliable half-maximal inhibitory concentration values (IC_50_) in the low µM range (Table 1, Fig. S1-S5). To further characterize the DNA damage response induced by the model colibactins, we evaluated recruitment of the p53-binding protein 1 (53BP1), a downstream effector of ATR kinase signaling.^*38*^ Of the suite of compounds evaluated, only **6** demonstrated significant levels of 53BP1 foci co-localized with γH2AX foci comparable to those induced by 1 µM etoposide or infection with *clb*-expressing strains at an MOI of 10 (Fig. 1, Fig. S6).^*20*^ Compounds **2** and **4** generated very weak 53BP1 signaling, with high micromolar concentrations demonstrating approximately 40% activity of the etoposide control (Fig. S1, S4). The coincidence of γH2AX and 53BP1 seen in select compounds supports activation of the eukaryotic DNA DSB repair pathways, without targeted cellular delivery or bacterial-human cell-to-cell contact.

**Table 1.** Half-maximal effective concentrations for γH2AX activation. Effect of compounds on γH2AX foci was quantified relative to 1 µM etoposide and graphed on a XY plot with a nonlinear regression curve fit. IC_50_ values were calculated unless the fit was ambiguous (n.d.0.

| Compound | Cell Line | IC<sub>50</sub> |
| --- | --- | --- |
| 2 | HeLa | n.d. |
|  | U2OS | n.d. |
| 3 | HeLa | 2.11 μM |
|  | U2OS | n.d. |
| 4 | HeLa | 3.15 μM |
| | U2OS | 2.87 μM |
| 5 | HeLa | 10.7 μM |
| | U2OS | 35.53 μM |
| 6 | HeLa | 65.68 μM |
| | U2OS | 1.28 μM |

Previously, **2** was shown to be a substrate for the resistance protein ClbS,^*27, 39*^ a cyclopropane hydrolase that catalyzes hydrolytic ring-opening of the cyclopropane.^*27*^ We showed that several products were spontaneously generated from the hydrolysis product including a proposed alkyl hydroperoxide. Both *N*-methylamide **2** and peroxide-containing hydrolytic intermediates possess a second electrophilic site (C4) in the lactam, which is proposed to account for the very weak DNA interstrand crosslinking activity previously observed for a truncated precolibactin *in vitro*^*21, 27*^ The potential role for this electrophilic reactivity *in vivo* is unknown.

Given the weak 53BP1 activation for **2**, we suspected that **2** and **3** might produce unstable crosslinks via the electrophilic lactam C4 that were not detected in our earlier study^*19*^. During these investigations, we noted that our synthetic colibactins were more stable at acidic pH (data not shown). We hypothesized that lowering the pH of the alkylation assay could increase the stability of the compounds in solution, resulting in stronger cross-linking activity. Indeed, reproducible formation of DNA interstrand crosslinks was observed when linearized pBR322 DNA was incubated with 10 or 1 μM of **2** or **3** at pH 5, as compared to assays conducted at pH 8 (Fig. 2A, Fig. S7-S8). We propose that decreased pH may enhance the stability of the model colibactin–DNA complexes. The degree of crosslinking induced by compound **3** was time-dependent, while the stability of DNA crosslinks induced by both **2** and **3** were decreased as the concentration of base arose during denaturing gel assays (Fig 2A, Fig. S7-S9). However, crosslinks resulting from natively expressing *clb*+ strains remain stable after denaturing with higher NaOH percentages (Fig. S11). The data indicate that colibactin analogs, which closely resemble known metabolites detected in cell culture, can crosslink DNA *in vitro*, although these crosslinks showed different stability compared to DNA crosslinked by native *clb*+ strains.

**Figure 2.**
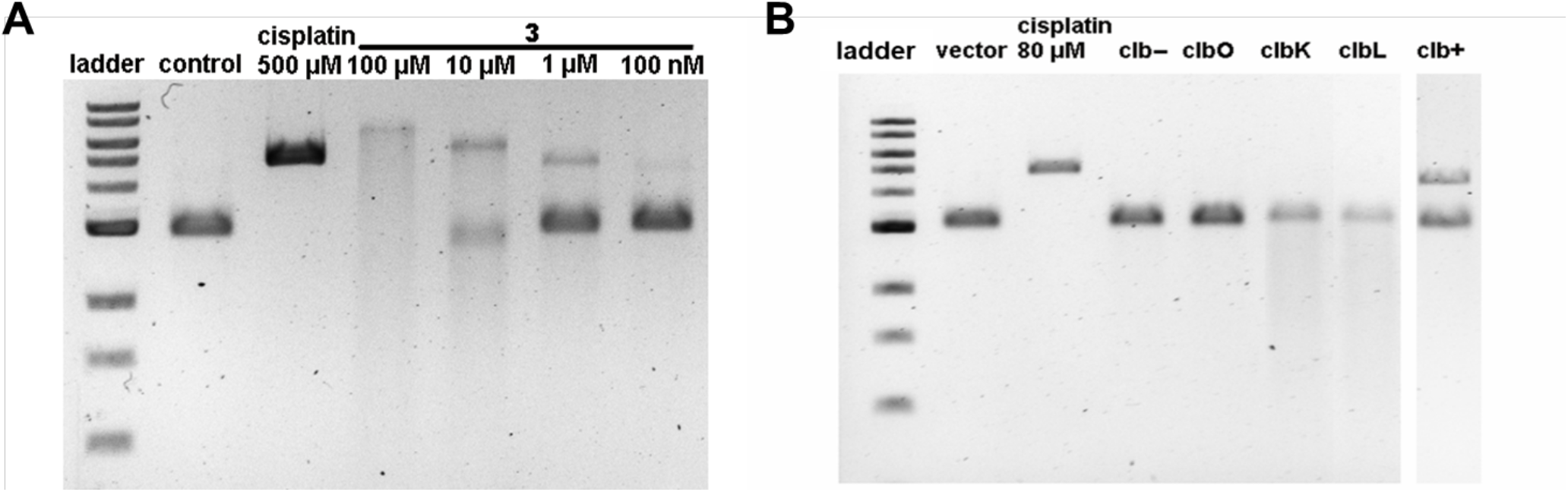
Model colibactins are able to both strongly alkylate and weakly crosslink DNA, compared to the full biosynthetic pathway capable of producing a major metabolite that strongly crosslinks DNA and minor pathway intermediates that alkylate DNA, as a result of evolutionarily encoded product diversification. (A). Incubation of Compound **3** with linearized pBR322 at pH 5 reveals clear crosslinks under mild denaturing conditions (50 mM/0.2% NaOH, on ice). (B). Incubation of linearized pBR322 with isogenic mutants of the *clb* pathway for 4 hours demonstrate varying levels of DNA alkylating activity. Denaturing gels were run at 50 mM (0.2%) NaOH.

Compounds **2**–**6** are approximately ten-fold less potent than etoposide in tissue culture, yet they induce comparable levels of DNA damage at nanomolar concentrations *in vitro*. We speculated that this discrepancy is due to issues of transport and membrane permeability and might be related to biosynthetic events not represented in the current model colibactins. The model colibactins employed in this study do not contain an αaminomalonate extender unit that is known to be incorporated by the PKS module of NRPS-PKS hybrid protein ClbK.^*23, 24, 30*^ While the genes dedicated to the synthesis and incorporation of this residue are essential for genotoxicity in the transient infection model, the only colibactin metabolite known to date that contains this motif has not been reported to harbor genotoxicity similar to fully functionalized native colibactin. This is in part due to the retention of the *N*-acyl-D-asparagine side chain, as well as possibly due to the missing biosynthetic additions provided by the largely uncharacterized enzymes ClbO and ClbL in the pathway. Though the exact functional requirement of the α-aminomalonate unit is as-of-yet unclear, it has been speculated that the basic amine in this residue may increase DNA affinity^*19, 24*^ and enhance cellular permeability. Indeed, a genetic screen of bleomycin-resistant S*accharomyces cerevisiae* revealed mutations in polyamine transporters as a key bleomycin-resistant determinant, implying that an amino-containing cationic cap could enhance nuclear delivery.^*40, 41*^

To further explore these functional consequences of the unaccounted *clb* enzymes, we evaluated the genotoxicity of pathway mutants both *in vitro* and in the transient infection model in comparison to our model colibactins. Active site point mutations were constructed in the PKS module of ClbK and in the peptidase ClbL based on protein sequence alignments to known ketosynthase and homologous amidase domains, as previously described.^*29*^ U2OS cells were transiently infected with DH10B *E. coli* carrying the full *clb* pathway, *clbK* point mutant (C167A), *clbL* point mutant (S179A), or empty BAC vector control, and γH2AX activation was analyzed by flow cytometry (Fig. S10). Both point mutants ablated the genotoxic effects of the pathway, demonstrating that these specific catalytic activities are essential for *clb+ E. coli* in inducing DSBs in cell culture.

To remove uncharacterized human cell trafficking aspects of the transient infection model, we next analyzed these strains for their ability to damage exogenously supplied DNA using denaturing gel assays (Fig. 2B). *E. coli* negative controls lacking the pathway (*clb-*) or fully deleted for the PKS *clbO* were inactive as expected. However, *E. coli* with the inactive PKS module in *clbK* or inactive peptidase *clbL* exhibited DNA damage, but not crosslinking. Because the complete pathway leads to stable DNA interstrand crosslinks, as described above and elsewhere^*20*^, relative to the model colibactins (Fig. S11), additional structural modifications afforded by the PKS module of ClbK, PKS ClbO, and amidase ClbL transform the potent DNA alkylators with weak-to-moderate crosslinking activities into a highly efficient DNA interstrand crosslinker. The activity of the *clbK* and *clbL* mutants *in vitro* is consistent with the extensive DNA alkylation and degradation that we have observed for model colibactins, which are analogs of metabolites produced by biosynthetic PKS module skipping of ClbK. Thus, while model colibactins do not fully reproduce the cellular phenotypes induced by fully functionalized colibactins, they do phenotypically recapitulate a subset of characterized on-pathway intermediates. This illustrates how product diversification evolutionarily encoded by the modular NRPS-PKS biosynthetic enzymes can generate functionally distinct products with varying modes of action. In addition, the alkylation activity for *clbK* and *clbL* mutants *in vitro*, but not in the transient infection model, again suggests that these uncharacterized enzymatic modifications enhance transport and cellular permeability.

The nature of colibactin transport is largely unknown; however, bacterial-human cell-to-cell contact is necessary for genotoxicity.^*9*^ This has largely puzzled researchers, given that expression of the *clb* locus does not depend on the presence of mammalian cells^*42*^ and outer membrane vesicles isolated from *clb*-expressing strains in some studies confer genotoxicity when applied,^*43*^ yet membranes separating *clb*-expressing bacteria from mammalian cells or thick adherent intestinal mucus layers abolishes genotoxicity.^*9, 44*^ To test whether the genotoxic effects of synthetic colibactins were eliminated by membrane separation, we treated U2OS cells with 50 μM **3** for 4 h in the presence or absence of separating 0.45 µm membranes, and γH2AX activation was analyzed by flow cytometry (Figure 3). While the effect was not drastic in comparison to the complete ablation of genotoxicity caused by the membrane in the presence of *clb+ E. coli*, the membrane did diminish the γH2AX response of **3** (Figure 3). To test whether the presence of bacteria affected the activity of **3**, we pretreated the membranes with *clb–* or *clb+* bacteria for 30 min before adding **3**. The γH2AX response was also diminished in this experiment. This suggests that diffusion of **3** across the membrane is slow, or potentially deleterious interactions with the membrane itself impede transport into cellular targets attenuating genotoxic activity.

**Figure 3.**
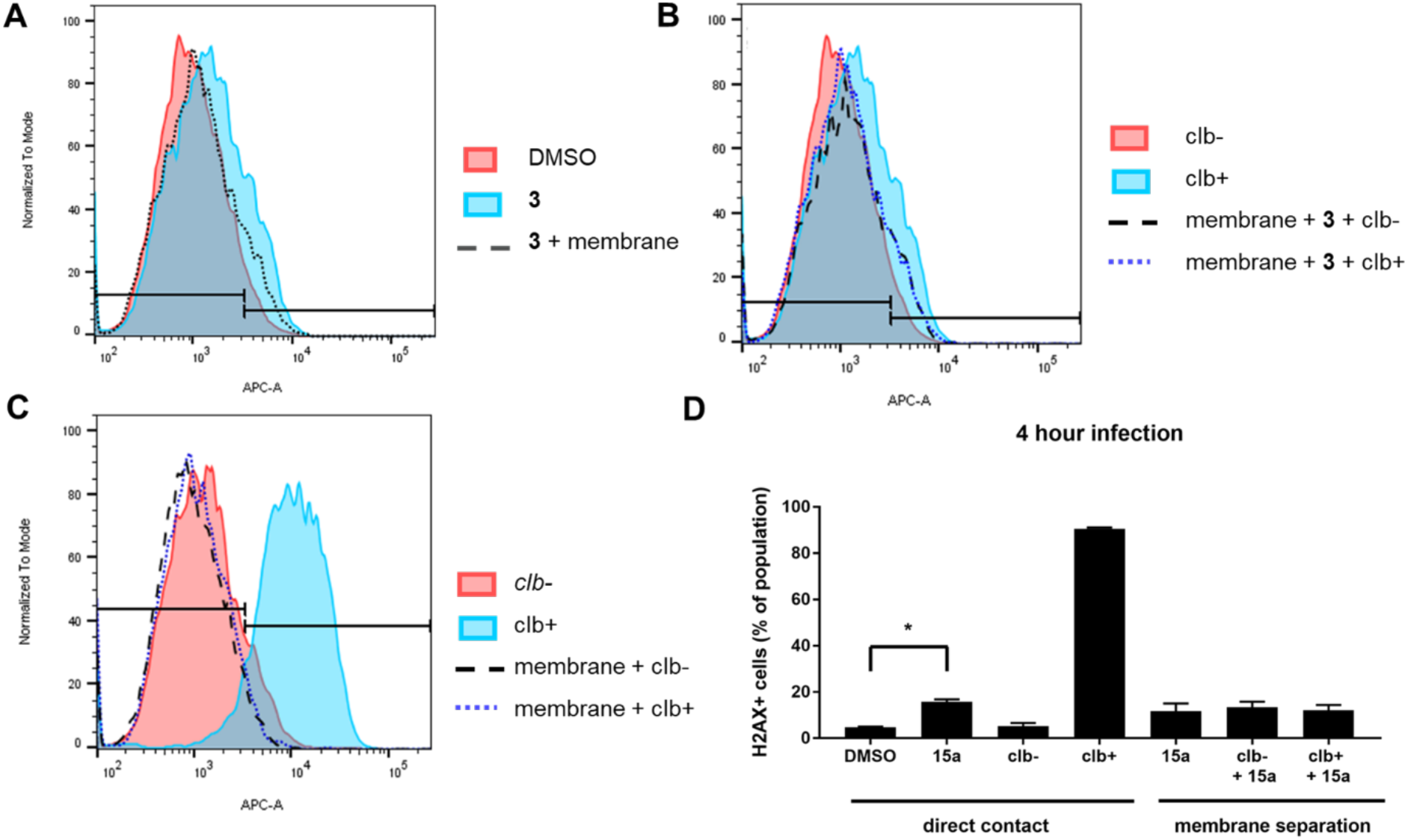
(A) 6.0 × 10^5^ U2OS cells were treated with 50 µM compound **3** in the presence or absence of a 0.45 µm membrane separating the bacteria from the U2OS cells for four hours. Cells were immediately fixed and stained for γH2AX and quantified by flow cytometry. (B) Membranes were pretreated with equal numbers of *clb-* or *clb+* bacteria grown to OD = 1.0 at MOI = 10 for 30 minutes prior to addition of **3** for four hours. (C) Native colibactins from live expressing *clb+* or control (*clb-*) cells applied with or without membranes to U2OS cells, and subsequently evaluated for DNA double stranded breaks. (D). Percentage of H2AX+ positive foci of each treatment condition, normalized to 5% H2AX+ for the negative control, DMSO vehicle. Results are shown as means and standard deviation (s.d.). *P* value = 0.0241, (* = < 0.05). *P* value obtained with an unpaired *t-* test with Welch’s correction.

Given that free model colibactins induce DNA double strand breaks in human cells, we evaluated whether ClbS could protect human cells when delivered extracellularly. We found that when purified ClbS was exogenously supplemented (at 1 µM in our studies), U2OS cells were protected from *clb+ E. coli*, as determined by quantification of γH2AX staining (Fig. 4; 10 µM ClbS studies are shown in Fig. S12). While this work was in progress, a similar observation was reported.^*20*^ Additionally, we had previously identified the active site Tyr residue Y55 in ClbS as critical for cyclopropane hydrolase activity.^*27*^ When 1 µM of the ClbS Y55F mutant was supplemented to U2OS cells transiently infected with *clb+ E. coli*, protective effects were not observed (Fig. 4). These data indicate that the previous protein biochemical studies using purified ClbS and model colibactins are consistent with the activities observed in cell cultures. These data also suggest that native colibactins can be intercepted by extracellularly supplemented ClbS. The results of these experiments also suggest that direct cell-to-cell contact might not be required for native colibactin activity. We propose that poor diffusion and chemical instability contribute to the observed bacteria-human cell-to-cell contact phenotype.

**Figure 4.**
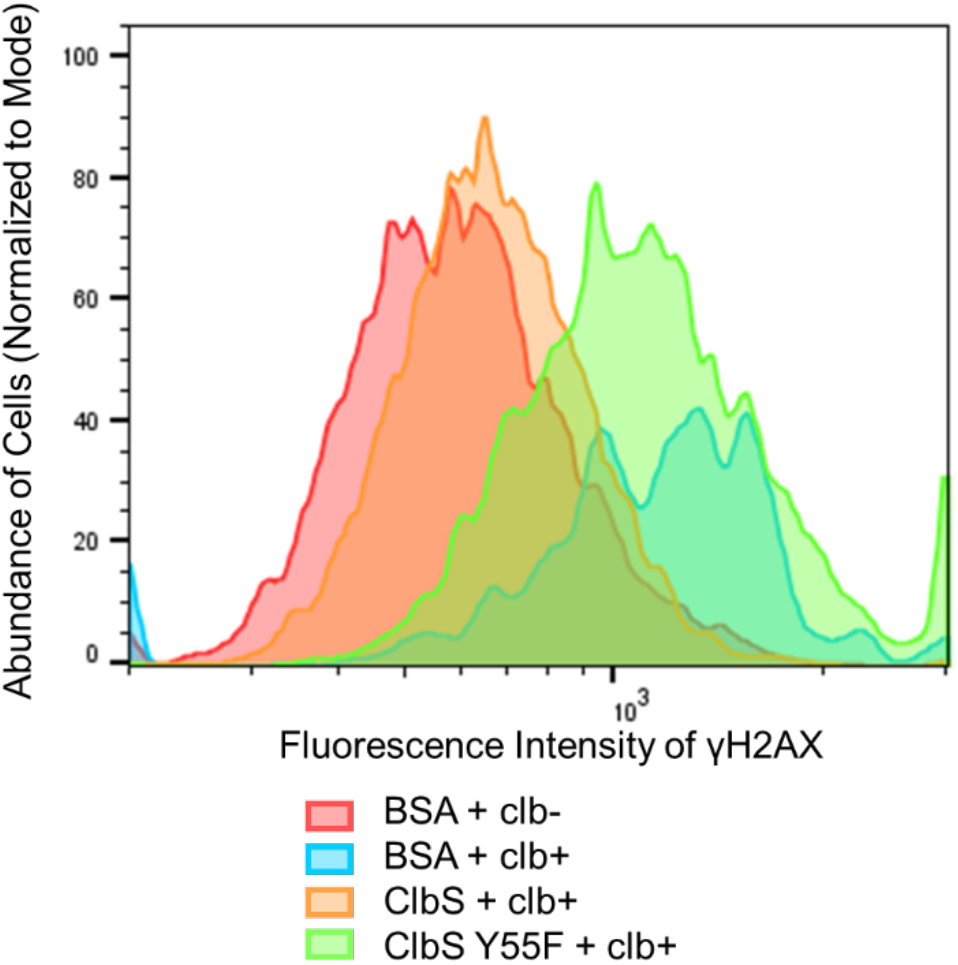
Protection from Genotoxicity by Supplementation of Extracellular ClbS. 6.0 × 10^5^ U2OS cells were transiently infected with *clb*-expressing bacteria at MOI = 10 for four hours. Media was replaced and 50 µg/mL gentamycin were added. Cell fixation and γH2AX antibody staining occurred 16 hours after infection to monitor downstream DSBs. γH2AX positive cells were quantified using flow cytometry.

In conclusion, we have demonstrated that model colibactin analogs can recapitulate cellular genotoxic phenotypes consistent with native *clb+ E. coli* infection by γH2AX and 53BP1 foci formation. In addition, we have shown that these compounds also reconstitute unstable DNA interstrand crosslinks *in vitro*. DNA interstrand crosslinking has recently been supported as the primary mode of action of the mature, final pathway product^*20*^. However, several modifications relative to the model colibactins, which are encoded in the PKS module of ClbK, PKS ClbO, and peptidase ClbL, are needed to recapitulate efficient stable crosslinks of the native colibactins produced by the full pathway. Our membrane studies, in combination with the ability of exogenous ClbS to intercept and neutralize native colibactins, call into question the absolute necessity of cell-to-cell contact for colibactin’s genotoxic action. Our work also provides important new insights toward resolving the remaining questions surrounding colibactin’s mature structure and mode of action.

## Methods

### Mammalian cell Lines and Reagents

U2OS and HeLa cell lines were obtained from American Type Culture Collection (ATCC). For microscopy, U2OS cells were cultured in McCoy’s 5A medium (Fisher Scientific) supplemented with 10% fetal bovine serum (FBS) (Life Technologies). For flow cytometry studies, U2OS cells were cultured in Dublecco’s Modified Eagle’s Medium (DMEM, Life Technologies)/ F12 with 25 mM HEPES and 5% FBS. HeLa cells were cultured in DMEM with 10% FBS. All cells were maintained at 37 °C with 5% CO2. Anti-bodies for immunofluorescence were purchased from Upstate (phospho-specific H2AX, 05–636), Novus Biologicals (53BP1, NB100–904SS), or Molecular Probes (Alexa Fluor 488-conjugated goat anti-mouse immunoglobulin G (IgG) and Alexa Fluor 647-conjugated goat anti-rabbit IgG). Antibodies for flow cytometry quantification were purchased from Cell Signaling (P-histone H2AX S139 rabbit) and Life Technologies (Alexa Fluor 647, REF A21245).

### Bacterial Strains and Growth Conditions

The *E. coli* strains used in this study were DH10B containing the colibactin gene cluster on pBeloBAC11 (*clb+*) or empty vector alone (*clb-*)^*9*^. The *clbL, clbK*, and *clbO* mutants were constructed as previously described^*29*^. Bacteria were grown overnight in LB and diluted into DMEM F12 15 mM HEPES with chloramphenicol (25 µg/mL) for assays. Standard growth conditions were 37 °C with 250-rpm agitation.

### Synthesis of Colibactin Analogs

Compounds were prepared as previously described.^*19*^

### Immunofluorescence Assay

Cells were seeded at 2500 (HeLa) or 5000 (U2OS) cells per well to achieve total well volumes of 20 μL in 384-well plates (black with optically clear bottom, Greiner Bio One 781091) using a Thermo Combidrop liquid dispenser. Cells were grown for 24 h, followed by the addition of test compounds using Echo acoustic liquid handler (Labcyte). For each tested drug concentration, 20 nL aliquot of the 1000× stock was added to 20 μL of cells to provide final DMSO concentration of 0.1%. Each plate contained 16 negative vehicle control wells (0.1% DMSO) and 16 positive control wells (1 μM etoposide). The cells were incubated with the compounds for 4 h and then fixed and subjected to immunofluorescence.

### Immunofluorescence

Cells were fixed with 4% paraformaldehyde (PFA) (Electron Microscopy Sciences) in the presence of 0.02% Triton X-100 at room temperature for 20 min and then incubated in permeabilization/blocking solution (10% FBS, 0.5% Triton X-100 in phosphate-buffered saline (PBS)) at room temperature for 1 h. Primary antibodies were diluted 1:500 in permeabilization/blocking solution and used to stain cells at 4 °C overnight. Cells were imaged using the InCell 2200 Imaging System (GE Corporation), and analyzed using InCell Analyzer software (GE Corporation) to quantify the number of γH2AX and 53BP1 foci. The effect of the test compounds was normalized to the mean of positive control wells (1 µM etoposide, set as 100% effect) and the mean of negative control well (0.1% DMSO, set as 0% effect).

### DNA crosslinking assay

The 4,163 bp plasmid pBR322 was purchased from NEB and linearized with 5U/μg EcoRI (NEB). The cut plasmid was purified using PCR clean kit (NEB) and eluted into 10 mM Tris pH 8.0. For each reaction with synthetic colibactin **2–6**, 200 ng of linearized DNA (31 μM base pairs) was incubated with compound in a 20 μL total volume. Compounds were diluted in DMSO such that each reaction consisted of a fixed 5% DMSO concentration. Reactions were conducted in 10 mM Tris EDTA buffer pH 8.0 or in 10 mM sodium citrate pH 5.0 as labeled on the figure. Reactions proceeded for 3 hours at 37 ºC, unless otherwise noticed. For each reaction with bacteria, 600 ng linearized plasmid DNA was added to 150 μL of DMEM/F12 15 mM Hepes (Invitrogen) inoculated with 6 ×10^6^ bacteria pre-grown to exponential phase in the DMEM/F12 15 mM Hepes (Invitrogen) media. The DNA-bacteria mix were incubated at 37 ºC for 4 hours, and the bacteria were then pelleted. The DNA was isolated from the supernatant using PCR clean kit (NEB) and quantified using nanodrop. The DNA concentration was adjusted to 10 ng/ μL using water and stored in −20 ºC until ready for gel analysis. Pure methyl methanesulfonate (MMS) (Alfa Aesar) and cisplatin (Biovision) stock solutions were diluted into DMSO immediately prior to used. As controls, 200 ng of DNA was treated with 80 μM of cisplatin (Biovision) in 10 mM sodium citrate pH 5 buffer with 5% final DMSO concentration. The DNA was immediately tested with gel electrophoresis after incubation.

### DNA gel electrophoresis

For each DNA sample, the concentration was pre-adjusted to 10 ng/ μL. 4 μL (40 ng) of DNA was taken out and mixed with 1.5 μL of 6× purple gel loading dye; no SDS (NEB) for non-denatured gel. For denatured gels, 5 μL (50 ng) of DNA was taken out each time and separately mixed with 15 μL of 0.2% denaturing buffer (0.27% sodium hydroxide, 10% glycerol, 0.013% bromophenol blue), 0.4% denaturing buffer (0.53% sodium hydroxide, 10% glycerol, 0.013% bromophenol blue), or 1% denaturing buffer (1.33% sodium hydroxide, 10% glycerol, 0.013% bromophenol blue) on an ice bath. The mixed DNA samples were denatured in 4 ºC for 10 min and immediately loaded onto 1% agarose Tris Borate EDTA (TBE) gels for 1.5 hour at 90 V. The gel was post stained with SybrGold (Thermo Fisher) for 2 hours.

### Membrane Transport Assay

6.0 × 10^5^ U2OS cells were seeded in 6-well plates and treated with 50 µM compound **3** diluted in 300 µl suspended on the surface of a 0.45 µm 30 mm-diameter membrane (Milipore) for four hours. DH10B *clb-* or *clb+* cells at OD600 = 1.0 were diluted into 300 µl DMEM at MOI of 10 and suspended on the surface of the 0.45 µm membrane to separate bacteria from the U2OS cells.

### ClbS Protection Assay

Expression and purification of ClbS was performed as previously described.^*27*^ Briefly, 1 L of BL21 cells carrying either pET28-ClbS-His or pET28-ClbSY55C were induced with 1 mM isopropyl β-D-thiogalactopyranoside (IPTG) and grown overnight at 25°C. Bacteria were lysed in lysis buffer (50 mM NaH2PO4, 300 mM NaCl, 10 mM imidazole, pH 8, 1 mg/mL of lysozyme freshly added) and purified on a 2 mL bed volume of HisPur Ni-NTA resin (Thermo Scientific). Proteins were eluted into 250 mM imidazole, 100 mM Tris, 300 mM NaCl, 10% glyercol (pH 8). Purified proteins were separated over a 15% SDS-PAGE gel to verify size and relative purity, and then buffer exchanged into 50 mM potassium phosphate buffer (pH 8) using PD-10 Desalting Columns (GE). 6.0 × 105 U2OS were infected with either *clb-* or *clb+* bacteria at OD600 = 1.0, MOI 10 with either 1 µM or 10 µM ClbS/ClbSY55C or BSA and incubated at 37 °C for 4 hours.

### Quantification of H2AX

Cells were immediately fixed and stained for γ-H2AX antibody and quantified by flow cytometry. Cells were fixed with 4% paraformaldehyde in PBS for 10 minutes, permeabilized with 90% ice cold methanol for 30 minutes on ice, and blocked in PBS with 1 mM CaCl2, 1 mM MgCl2, and 10% FBS for 10 minutes. Primary antibodies were diluted 1:100 in blocking solution and incubated overnight at 4°C. Secondary antibodies were diluted 1:1000 in blocking solution and incubated at room temperature for 2 hours. Cells were washed and resuspended into PBS before analysis on a FACSAria II (BD).

## Associated Content

**Supporting Information**

Figures S1-S12 (PDF).

**Notes**

The authors declare no competing financial interests.

## Acknowledgment

Financial support from the National Institutes of Health (1DP2-CA186575 to J.M.C., R01GM110506 to S.B.H., and R01CA215553 to S.B.H & J.M.C), the Burroughs Wellcome Fund (1016720 to J.M.C.), the Camilee & Henry Dreyfus Foundation (TC-17–011 to J.M.C.), and Yale University is gratefully acknowledged. E.E.S. was supported by the National Science Foundation Graduate Research Fellowships Program. A.R.H. was supported by a Charles H. Revson Foundation Senior Fellowship.

---

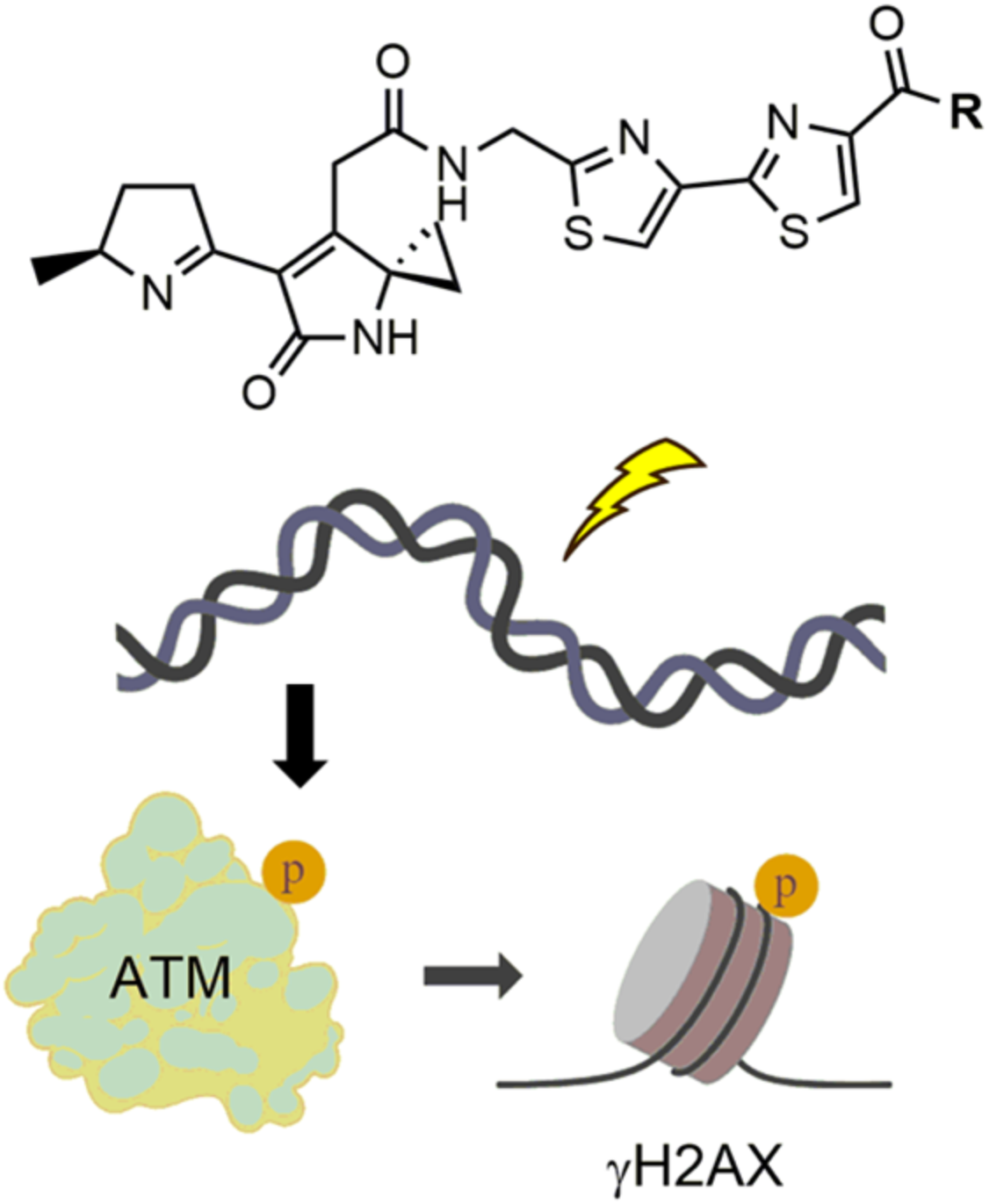

## References

1. Winter, S. E., Lopez, C.A., Baumler, A.J. (2013) The dynamic of gut-associated microbial comunities during inflammation, EMBO Reports 14, 319–327.

2. Garrett, W. S. (2015) Cancer and the microbiota, Science 348, 80–86.

3. Gilbert, J. A., Quinn, R.A., Debelius, J., Xu, Z.Z., Morton, J., Garg, N., Jansson, J.K., Dorrestein, P.C., Knight, R. (2016) Microbiome-wide association studies link dynamic microbial consortia to disease, Nature Chemical Biology 535, 94–103.

4. Balskus, E. P. (2015) Colibactin: understanding an elusive gut bacterial genotoxin, Natural Product Reports 32, 1534–1540

5. Trautman, E. P., Crawford, J.M. (2016) Linking Biosynthetic Gene Clusters to their metabolites via Pathway Targeted Molecular Networking, Current Topics in Medicinal Chemistry 16, 1705–1716.

6. Taieb, F., Petit, C., Nougayrede, J.P., Oswald, E. (2016) The Enterobacterial Genotoxins: Cytolethal Distending Toxin and Colibactin EcoSal Plus doi:10.1128/ecosalplus.ESP-0008-2016.

7. Healy, A. R., Herzon, S.B. (2017) Molecular Basis of Gut Microbiome-Associated Colorectal Cancer: A Synthetic Perspective, J. Am. Chem. Soc. 139, 14817–14824.

8. Fais, T., Delmas, J., Barnich, N., Bonnet, R., Dalmasso, G. (2018) Colibactin: More Than a New Bacterial Toxin, Toxins 10.

9. Nougayrede, J.-P., Homburg, S., Taieb, F., Boury, M., Brzuszkiewicz, E., Gottschalk, G., Buchrieser, C., Hacker, J., Dobrindt, U., and Oswald, E. (2006) Escherichia coli induces DNA double-strand breaks in eukaryotic cells, Science 313, 848–851.

10. Johnson, J. R., Johnston, B., Kuskowski, M.A., Nougayrede, J.P., Oswald, E. (2008) Molecular Epidemiology and Phylogenetic Distribution of the Escherichia coli pks Genomic Island, Journal of Clinical Microbiology 46, 3906–3911.

11. Putze, J., Hennequin, C., Nougayrede, J.P., Zhang, W., Homburg, S., Karch, H., Bringer, M.A., Fayolle, C., Carniel, E., Rabsch, W., Oelschlaeger, T.A., Oswald, E., Forestier, C., Hacker, J., and Dobrindt, U. (2009) Genetic Structure and Distribution of the Colibactin Genomic Island among Members of the Family Enterobacteriaceae, Infection and Immunity 77, 4696–4703.

12. Arthur, J., Perez-Chanona, E., Muhlbauer, M., Tomkovich, S., Uronis, J., Fan, T., Campbell, B., Abujamel, T., Dogan, B., Rogers, A., Rhodes, J., Stintzi, A., Simpson, K., Hansen, J., Keku, T., Fodor, A., and Jobin, C. (2012) Intestinal inflammation targets cancer-inducing activity of the microbiota, Science 338, 120–123.

13. Buc, E., Dubois, D., Sauvanet, P., Raisch, J., Delmas, J., Darfeuille-Michaud, A., Pezet, D., and Bonnet, R. (2013) High prevalence of mucosa-assoicated E.coli producing cyclomodulin and genotoxin in colon cancer, PLOS ONE 8, e56964.

14. Dejea, C. M., Fathi, P., Craig, J.M., Boleij, A., Taddese, R., Geis, A.L., Wu, X., DeStefano Shields, C.E., Hechenbleikner, E.M., Huso, D.L., Anders, R.A., Giardiello, F.M., Wick, E.C., Wang, H., Wu, S., Pardoll, D.Rm., Housseau, F., Sears, C.L. (2018) Patients with Familian Adenomatous Polyposis Harbor Colonic biofilms containing tumorigenic bacteria, Science 359, 592–597.

15. Cuevas-Ramos, G., Petit, C. R., Marcq, I., Boury, M., Oswald, E., and Nougayrede, J.-P. (2010) Escherichia coli induces DNA damage in vivo and triggers genomic instability in mammalian cells, Proc. Natl. Acad. Sci. U.S.A. 107, 11537–11542.

16. Secher, T., Samba-Louaka, A., Oswald, E., Nougayrede, J.P. (2013) Escherichia coli Producing Colibactin Triggers Premature and Transmissible Senescence in Mammalian Cells, PLOS ONE 8, e77157.

17. Bonnet, M.,, Buc, E., Sauvanet, P., Darcha, Cl., Dubois, D., Pereria, B., Dechelotte, P., Bonnet, R., Pezet, D., Darfeuille-Michaud, A. (2014) Colonization of the Human Gut by E. coli and Colorectal Cancer Risk, Clinical Cancer Research 20, 859–867.

18. Tomkovich, S., Yang, Y., Winglee, K., Gauthier, J., Mühlbauer, M., Sun, X., Mohamadzadeh, M., Liu, X., Martin, P., Wang, G.P., Oswald, E., Fodor, A.A., Jobin, C. (2017) Locoregional Effects of Microbiota in a Preclinical Model of Colon Carcinogenesis, Cancer Res. 77, 2620–2632.

19. Healy, A. R., Nikolayevskiy, H., Patel, J. R., Crawford, J. M., and Herzon, S. B. (2016) A mechanistic model for colibactin-induced genotoxicity, J. Am. Chem. Soc. 138, 15563–15570.

20. Bossuet-Greif, N., Vignard, J., Taieb, F., Mirey, G., Dubois, D., Petit, C., Oswald, E., Nougayrede, J.P.. (2018) The Colibactin Genotoxin Generates DNA Interstrand Cross Links in Infected Cells, mBio 9, e02393–02317.

21. Vizcaino, M. I., and Crawford, J. M. (2015) The colibactin warhead crosslinks DNA, Nature Chemistry 7, 411–417.

22. Brotherton, C. A., and Balskus, E. P. (2013) A prodrug resistance mechanism is involved in colibactin biosynthesis and cytotoxicity, Journal of the American Chemical Society 135, 3359–3362.

23. Brachmann, A. O., Garcie, C., Wu, V., Martin, P., Ueoka, R., Oswald, E., and Piel, J. (2015) Colibactin biosynthesis and biological activity depend on the rare aminomalonyl polyketide precursor, Chem. Commun. 51, 13138–13141.

24. Zha, L., Wilson, M. R., Brotherton, C. A., and Balskus, E. P. (2016) Characterization of polyketide synthase machinery from the pks island facilitates isolation of a candidate precolibactin, ACS Chem Biol 11, 1287–1295.

25. Guntaka, N. S., Healy, A. R., Crawford, J. M., Herzon, S. B., and Bruner, S. D. (2017) Structure and functional analysis of ClbQ, an unusual intermediate-releasing thioesterase from the colibactin biosynthetic pathway, ACS Chem Biol 12, 2598–2608.

26. Zha, L., Jiang, Y., Henke, M. T., Wilson, M. R., Wang, J. X., Kelleher, N. L., and Balskus, E. P. (2017) Colibactin assembly line enzymes use S-adenosylmethionine to build a cyclopropane ring, Nature Chemical Biology 13, 1063–1065.

27. Tripathi, P., Shine, E.E., Healy, A.R., Kim, C.S., Herzon, S.B., Bruner, S.D., Crawford, J.M. (2017) ClbS is a cyclopropane hydrolase that confers colibactin resistance, J. Am. Chem. Soc. 139, 17719–17722.

28. Vizcaino, M. I., Engel, P., Trautman, E., and Crawford, J. M. (2014) Comparative metabolomics and structural characterizations illuminate colibactin pathway-dependent small molecules, Journal of the American Chemical Society 136, 9244–9247.

29. Trautman, E. P., Healy, A. R., Shine, E. E., Herzon, S. B., and Crawford, J. M. (2017) Domain-targeted metabolomics delineates the heterocycle assembly steps of colibactin biosynthesis, J. Am. Chem. Soc. 139, 4195–4201.

30. Li, Z. R., Li, J., Gu, J.P., Lai, J.Y.H., Duggan, B.M., Zhang, W.P., Li, Z.L., Li, Y.X., Tong, R.B., Xu, Y., Lin, D.H., Moore, B.S., Qian, P.Y. (2016) Divergent biosynthesis yields a cytotoxic aminomalonate-containing precolibactin, Nature Chemical Biology 12, 773–778.

31. Li, Z.-R., Li, Y., Lai, J. Y. H., Tang, J., Wang, B., Lu, L., Zhu, G., Wu, X., Xu, Y., and Qian, P.-Y. (2015) Critical intermediates reveal new biosynthetic events in the enigmatic colibactin pathway, Chem Biochem 16, 1715–1719.

32. Healy, A. R., Vizcaino, M. I., Crawford, J. M., and Herzon, S. B. (2016) Convergent and modular synthesis of candidate precolibactins. Structural revision of precolibactin A, J. Am. Chem. Soc. 138, 5426–5432.

33. Bian, X., Fu, J., Plaza, A., Herrmann, J., Pistorius, D., Stewart, A. F., Zhang, Y., and Muller, R. (2013) In vivo evidence for a prodrug activation mechanism during colibactin maturation, Chem Biochem 14, 1194–1197.

34. Mousa, J. J., Newsome, R. C., Yang, Y., Jobin, C., and Bruner, S. D. (2017) ClbM is a versatile, cation-promiscous MATE transporter found in the colibactin bioysnthetic gene cluster, Biochemical and Biophysical Research Communications 482, 1233–1239.

35. Cougnoux, A., Gibold, L., Robin, F., Dubois, D., Pradel, N., Darfeuille-Michaud, A., Dalmasso, G., Delmas, J., and Bonnet, R. (2012) Analysis of structure-fuction relationships in the colibactin-maturaing enzyme clbP, J. Mol. Biol 424, 203–214.

36. Dubois, D., Baron, O., Cougnoux, A., Delmas, J., Pradel, N., Boury, M., Bouchon, B., Bringer, M.-A., Nougayrede, J.-P., Oswald, E., and Bonnet, R. (2011) ClbP is a prototype of a peptidase subgroup invovled in biosynthesis of nonribosomal peptide, J. Biol. Chem. 286.

37. Rogakou, E. P., Pilch, D.R., Orr, A.H., Ivanova, V.S., Bonner, W.M. (1998) DNA double-stranded breaks induce histone H2AX phosphorylation on serine 139., J Biol Chem 273, 5858–5868.

38. Ciccia, A., Elledge, S.J. (2010) The DNA damage response: making it safe to play with knives., Molecular Cell 40, 179–204.

39. Bossuet-Greif, N., Dubois, D., Petit, C., Tronnet, S., Martin, P., Bonnet, R., Oswald, E., and Nougayrède, J.-P. (2016) Escherichia coli ClbS is a colibactin resistance protein, Molecular Microbiology 99, 897–908.

40. Chen, J., Stubbe, J. (2005) Bleomycins: Towards Better Therapeutics, Nature Reviews Cancer 5, 102–112.

41. Aouida, M., Page, N., Leduc, A., Peter, M., Ramotar, D. (2004) A genome-wide screen in Saccharomyces cerevisiae reveals altered transport as a mechanism of resistance to the anticancer drug bleomycin, Cancer Res. 64, 1102–1109.

42. Homburg, S., Oswald, E., Hacker, J., Dobrindt, U. (2007) Expression analysis of the colibactin gene cluster coding for a novel polyketide in Escherichia coli, FEMS Microbiol. Lett. 275, 255–262.

43. Canas, M.-A., Gimenez, R., Fabrega, M.-J., Toloza, L., Baldoma, L., Badia, J. (2016) Outer membrane vesicles from the probiotic Escherichia coli Nissle 1917 and the Commensal ECOR12 enter intestinal epthileal cells via clathrin-dependent endocytosis and elicit differential effects on DNA damage, PLOS ONE 11, e0160374.

44. Reuter, C., Alzheimer M., Walles, H., Oelschlaeger, TA.. (2018) An adherent mucus layer attenuates the genotoxic effects of colibactin, Cell Microbiol 20, https://doi.org/10.1111/cmi.12812.

